# First clinical validation of whole genome screening on standard trophectoderm biopsies of preimplantation embryos

**DOI:** 10.1101/2022.04.14.488421

**Authors:** Yuntao Xia, Willy Chertman, Dhruva Chandramohan, Maria Katz, Elan Bechor, Ben Podgursky, Michael Hoxie, Qinnan Zhang, Jessica Kang, Edwina Blue, Justin Chen, Justin Schleede, Nathan Slotnick, Xiaoli Du, Jonathan Kort, Robert Boostanfar, Eric Urcia, Barry Behr, Noor Siddiqui

**Affiliations:** Orchid Health, Palo Alto, CA, 94301; Department of Obstetrics & Gynecology - Reproductive Endocrinology and Infertility, Stanford University, Sunnyvale, CA, 94087; RMA Northern California, San Francisco, CA, 94105; HRC Fertility-Encino, Encino, CA 91436

**Keywords:** Preimplantation genetic testing, Whole genome screening, In vitro fertilization, Next generation sequencing, Clinical validation

## Abstract

Whole Genome Sequencing (WGS) is used in healthcare and in the clinic, with the notable exception of preimplantation genetic testing (PGT). In PGT, only a few cells are available for sequencing, requiring DNA amplification which reduces data quality, sequence fidelity and sharply limits subsequent clinical impact. Here we demonstrate the first clinical validation of WGS on embryo biopsies using our lab development protocol, opening the door to broad use of WGS in fertility. We find that amplified DNA with comparable sensitivity and specificity to genomic DNA when performing whole genome sequencing assays. DNA amplification on cell lines and donated human embryos had an amplification success rate of >99.9% and 98.2% respectively and accuracy on both was >99.9% on aneuploidy status. GIAB samples (Genome in the Bottle reference NA12878) showed that our amplified DNA was broadly comparable to genomic DNA (99.99% accuracy, 99.99% specificity, 98.0% sensitivity and 98.1% precision). Using our assay, we were able to call variants, detect mitochondrial heteroplasmy, perform high precision screening without access to parental genomes, detect compound heterozygous variants, and score microdups/dels and uniparental disomies (to reduce risk of diseases such as DiGeorge syndrome and Prader-Willi syndrome). Our clinical study suggests that the full spectrum of traditional clinical genome bioinformatics, so far reserved to large samples, can now be performed on embryos before implantation.

## Introduction

Preimplantation genetic testing (PGT) reduces the number of cycles needed to achieve a successful pregnancy by improving fetal viability rates per embryo transfer(1). However, current methods can only assess ploidy status or screen for specific mendelian variants for which parents are known to be carriers, in which case DNA samples from both parents are required. However, for cases where donor eggs or donor sperm are used, it is difficult or impossible to gain access to donor DNA to evaluate carrier status(2). In addition, small copy number variations (CNV) such as DiGeorge are not screened in regular PGT-A because the deletion size falls below the sensitivity threshold (∼2Mb). Uniparental disomy (UPD) (such as Prader-Willi and Angelman) is also not regularly screened in PGT-A as low-pass next generation sequencing (NGS) data cannot accurately capture beta-allele frequency. To our knowledge, no other clinical whole genome sequencing is currently performed due to poor coverage and high allelic dropout rates(3). Here, we present a validation study on our latest whole-genome screening assay on embryos. We sought to measure the performance of this new assay against the current clinical standard in ploidy analysis, variant detection via WGS, and screening for mitochondrial genetic disorders, microduplications/deletions, and UPD from a single biopsy.

## Materials and Methods

### Sample collection

Cell lines were obtained from Coriell. Embryo samples were all donated to research and provided from the HRC Fertility Center (wIRB #20215134). All embryos were created using Intracytoplasmic sperm injection (ICSI). Embryos from day 5/6/7 were biopsied following standard SOP from clinics. No special SOP is needed. Each biopsy contained approximately 5 cells and was collected in a 200ul PCR tube with 3ul of cell buffer. Clinical case study was done under wIRB # 2022264.

### Next generation sequencing

Samples were processed in the Orchid lab (certified: CAP # 9234146 & CLIA # 34D2260214). DNA or biopsies were amplified using lab-developed protocol. DNA sizes after WGA were first confirmed by running 1-2% Agarose E-Gel (Invitrogen). After size determination, 250-500ng of DNA was used for library preparation with KAPA HyperPlus kit per manufacturer’s instructions. Dual Index UMI adapters (Integrated DNA Technologies) were used in the ligation. Library concentration was quantified using the Qubit 4 dsDNA HS. Library sizes were measured through Agilent 4150 Tapestation Genomic ScreenTape assay (Agilent Technologies). Sequencing runs were performed on a MiniSeq for low-pass aneuploidy screening and NovaSeq6000 for 30X WGS per manufacturer’s instructions.

### Genomic Data Analyses

NGS data were processed via a combination of Gencove Sentieon-based human-genome pipeline and an in-house pipeline following GATK best practices(4). Gvcfs were generated for each sample and jointly called as part of a larger cohort. GATK VariantRecalibrator (VQSR) was run on the resulting callset. Mitochondrial DNA was analyzed using GATK without VQSR. Sensitivity, precision, etc were calculated through RTG packages. NxClinical and Ginkgo were used for CNV calling using default parameters as the package is designed for amplified DNA. All data were aligned to GRCh37 with a bin size of 500 kbp. Embryos were excluded from the analysis based on the following criteria: 1) the whole embryo was deemed not viable or, 2) both the whole embryo and biopsies were inconsistent with previous PGT-A results.

## Results

### Validation on aneuploidy screening via cell lines and human embryos

Ploidy analysis was initially validated by assessing the accuracy of aneuploidy screening. A total of 73 samples from Coriell cell lines were used first to validate our aneuploidy screening (Figure S1). These cell lines covered common karyotypes seen in PGT-A, including euploid males and females and autosomal and sex chromosome aneuploidy. We diluted the DNA to ∼20-50 pg (picogram), which is equivalent to the amount of DNA in embryo trophectoderm biopsies. Whole-genome amplification was then performed, followed by low pass sequencing (∼0.01 - 0.05X) for aneuploidy evaluation with a bin size of 500 kbp on the genome in the analysis. Overall, we achieved >99.9% successful amplification and >99.9% accuracy (Figure S1B). Notably, we obtained >1500 ng of DNA for all samples, which is higher and more robust than most WGA approaches.

Next, we extended our PGT-A validation on donated human embryos with a larger variety of karyotypes (Figure 1, Table S1). We obtained 2-3 samples per embryo, on which we performed WGA and ploidy analysis. We first evaluated the consistency of these samples by comparing our CNV analysis to the previous clinical PGT-A reports. Among the samples, we observed a mosaic embryo with two samples being deletion of the short arm of chromosome 2 (−2p) and one sample being monosomy 2 (−2) (#55-57); and a mosaic embryo with one sample being mosaic trisomy 19 (+19) and one sample being trisomy 19 (+19). (#105-106). Considering the previous PGT-A results reported the same chromosome and the unknown accuracy of the original 2017 PGT-A testing technology, our PGT-A results are acceptably consistent. Two samples (marked indeterminate, Table S1) have more than 8% mitochondrial reads in the NGS data, negatively impacting our CNV calling; however, as high levels of mitochondrial reads often suggest embryo viability issues, the clinical impact of processing samples contaminated by high levels of mitochondrial DNA may be limited(5). Viability is generally higher with fr7esh embryos compared to banked embryos because banked embryos suffer from multiple freeze-thaw cycles and additional rounds of biopsies. Regardless, we achieved >99.9% accuracy overall with 1.8% indeterminate among 110 embryo samples.

**Figure 1.**
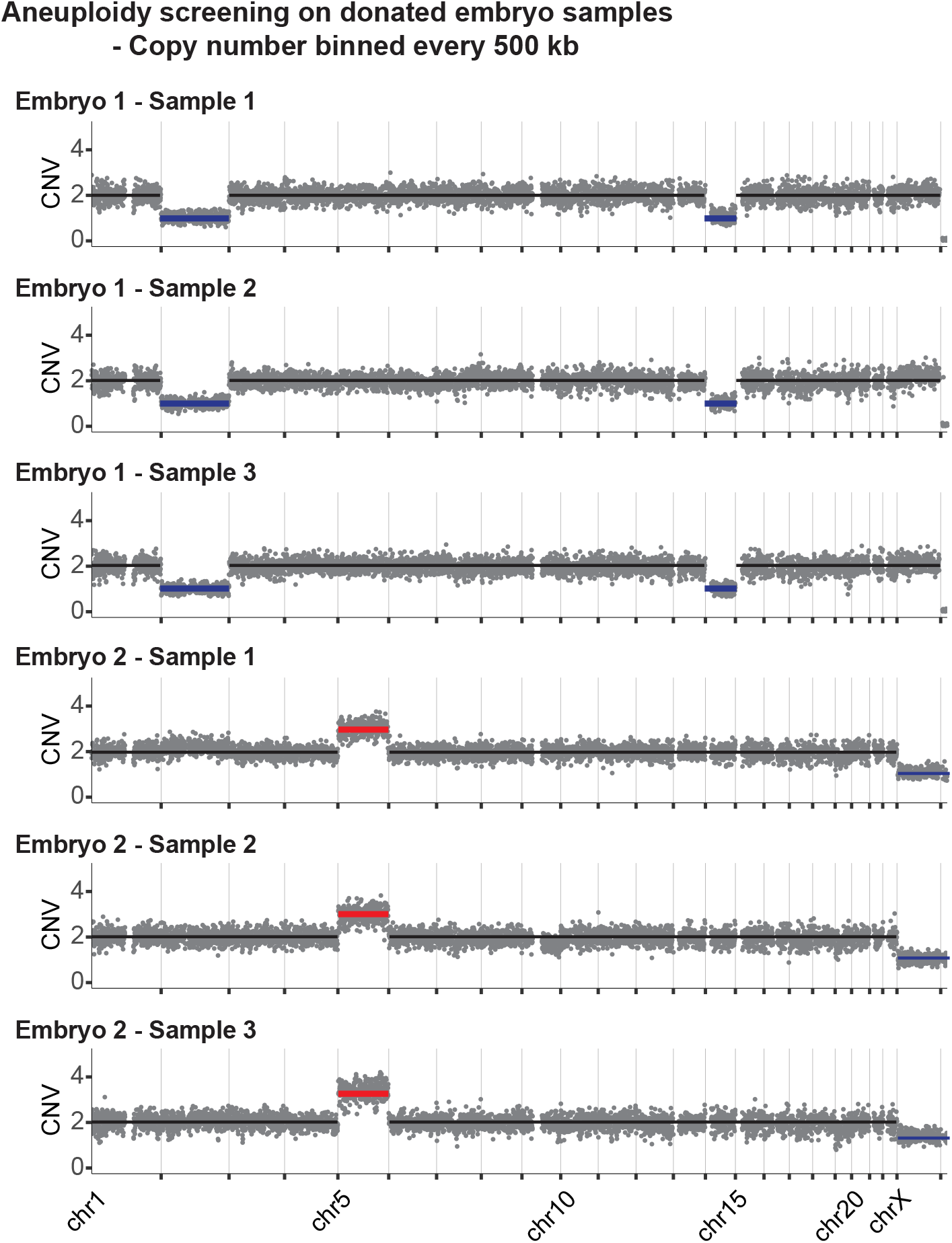
Summary of PGT-A on embryo samples. A) Examples of copy number results on embryo samples.

Low-level mosaicism in embryos has been reported to be compatible with a successful pregnancy and live birth, especially when the aneuploidy mosaicism level is less than 40%(6). Therefore, our algorithm was designed to capture mosaicism above 30%. We performed a dilution series from 50% to 10% mosaicism, generated by mixing aneuploid (47, XY, +18) and euploid (46, XX) DNA. As seen in Figure S2B-Batch1, mosaic samples above 30% could be visualized and were captured successfully by the algorithm. To confirm the results, we performed another mosaicism dilution using a mix of NA09288 (47, XY, +9) and euploid (46, XX) DNA and found the same result (Figure S2B-Batch2). Additionally, 24 cell line mixtures with 30-35% aneuploidy mosaicism were tested and 100% of them were successfully marked during analysis (Figure S2B-Batch3).

### Validation of whole genome sequencing using cell lines and human embryos

Although multiple clinical labs have demonstrated good performance in PGT-A(7), to our knowledge none have attempted validation on PGT-whole genome sequencing. DNA derived from whole genome amplification to date has poor quality, and lowered sequence fidelity when compared to genomic DNA and has not been used for embryo screening under clinical conditions(8). Our results using lab-developed WGA protocol show that our assay achieves comparable sensitivity and specificity to genomic DNA.

We used the NIST Genome in a Bottle – NA12878 and its variants as the data analysis reference. Both genomic DNA and amplified DNA from NA12878 were analyzed and compared to benchmark calls from GIAB (NIST v4.2.1). As shown in Table S2 and Table S3, accuracy and specificity at whole genome level are 99.99% for both amplified DNA and genomic DNA. We observed an average of 99.8% genomic coverage in amplified DNA and the same in gDNA (Table S2 and Table S3). The precision of amplified DNA and gDNA are 98.1% and 97.8%, respectively, suggesting our amplified DNA does not generate more false positive calls than gDNA (Table S2). Moreover, sensitivity reaches 98.0% in amplified DNA as compared to 98.9% in gDNA, indicating false negative calls in data are similar in magnitude as well (Table S2). Therefore, our WGS on amplified DNA was validated against the variant list from NIST and its sequencing outcome is comparable to gDNA.

To validate performance on biopsies of clinical embryos, we compared the results of the biopsies to their corresponding embryos. Using the whole embryo as the reference, we observed an average of 99.6% genomic coverage, 99.9% accuracy, 99.9% specificity, 98.0% precision, 98.1% sensitivity (Table 1 & Table S4). The demonstrated concordance allows us to treat WGS of an embryo biopsy as an accurate reflection of the entire embryo’s genome.

**Table 1.** Comparison of biopsies and their embryos in terms of genomic coverage, total SNV called, accuracy, specificity, precision, and sensitivity. The whole embryos were used as references.

| Sample | Genomic Coverage | Total SNV | Accuracy | Specificity | Precision | Sensitivity |
| --- | --- | --- | --- | --- | --- | --- |
| 1 | 99.7% | 3343804 | 99.995% | 99.997% | 97.8% | 98.4% |
| 2 | 99.6% | 3327631 | 99.995% | 99.997% | 98.0% | 98.1% |
| 3 | 99.6% | 3312139 | 99.995% | 99.997% | 97.9% | 98.1% |
| 4 | 99.4% | 3279468 | 99.993% | 99.997% | 97.8% | 97.1% |
| 5 | 99.6% | 3404155 | 99.995% | 99.998% | 98.4% | 98.1% |
| 6 | 99.7% | 3414277 | 99.996% | 99.998% | 98.4% | 98.4% |
| 7 | 99.6% | 3360682 | 99.995% | 99.998% | 98.4% | 98.1% |
| 8 | 99.6% | 3163591 | 99.995% | 99.998% | 98.2% | 98.2% |
| 9 | 99.6% | 3160113 | 99.995% | 99.998% | 98.3% | 98.2% |
| 10 | 99.6% | 3304310 | 99.994% | 99.996% | 97.2% | 98.0% |
| 11 | 99.7% | 3301716 | 99.994% | 99.997% | 97.5% | 98.2% |
| 12 | 99.7% | 3368834 | 99.996% | 99.998% | 98.6% | 98.2% |
| 13 | 99.7% | 3340483 | 99.995% | 99.998% | 98.8% | 97.5% |
| 14 | 99.6% | 3270893 | 99.995% | 99.997% | 97.5% | 98.4% |
| 15 | 99.5% | 3262566 | 99.995% | 99.997% | 97.6% | 98.3% |
| 16 | 99.6% | 3274334 | 99.995% | 99.997% | 97.8% | 98.2% |
| 17 | 99.6% | 3275227 | 99.995% | 99.997% | 97.7% | 98.2% |
| 18 | 99.6% | 3254447 | 99.995% | 99.998% | 98.1% | 98.4% |
| 19 | 99.4% | 3234717 | 99.995% | 99.998% | 98.2% | 97.9% |
| 20 | 99.7% | 3274002 | 99.995% | 99.997% | 97.9% | 98.5% |
| 21 | 99.5% | 3257361 | 99.995% | 99.998% | 98.1% | 98.2% |
| 22 | 99.6% | 3270572 | 99.995% | 99.998% | 98.2% | 97.9% |
| 23 | 99.7% | 3284833 | 99.995% | 99.998% | 98.2% | 98.3% |
| 24 | 99.6% | 3237713 | 99.993% | 99.997% | 97.9% | 97.0% |
| 25 | 99.7% | 3382545 | 99.996% | 99.998% | 98.3% | 98.6% |
| **Average** | **99.6%** | **3294417** | **99.995%** | **99.997%** | **98.0%** | **98.1%** |

### Validation on mitochondrial DNA

A high proportion of pathogenic heteroplasmy has been correlated to mitochondrial diseases including early onset metabolic and degenerative diseases. Our validation on mitochondrial DNA focused on mitochondrial genome coverage, sensitivity, and precision by comparing biopsies to their embryos. Setting 15% as our reportable heteroplasmy threshold, we observed an average of 100% mitochondria DNA coverage, >99.9% accuracy, >99.9% specificity, 99.0% sensitivity and >99.9% precision (Table S5). Since mitochondrial DNA is known to be exclusively inherited from the mother, our assay has the potential to be used to perform maternity confirmation.

### Detection of specific variants through WGS without parental samples

We performed WGS on biopsies from embryos that previously underwent PGT-M, as well as on Coriell cell lines that carry specific pathogenic variants. Following our protocol with 30X sequencing, all variants detected by previous PGT-M assays were detected successfully in our WGS results without the use of parental genomes (Table 2). To further confirm our ability to capture pathogenic variants in WGS, Coriell cell lines were selected to represent common genetic disorders. Amplified DNA from picogram-level of Coriell cell line followed by 30X sequencing further confirmed our PGT-WGS assay has ability to detect all reported pathogenic variants regardless of the number of variants in one sample (Table 3).

**Table 2.** Variants in embryo samples that underwent PGT-M previously. PGT-WGS captured all reported variants in positive samples and was clean on negative samples.

| Family | Sample | Previous PGT-M Reports | Gene | Disease | Detected in Orchid WGS? |
| --- | --- | --- | --- | --- | --- |
| Family 1 | Sample 1 | c.2279G>A | TGM1 | Lamellar Ichthyosis | Yes |
|  | Sample 2 | c.2279G>A | TGM1 | Lamellar Ichthyosis | Yes |
|  | Sample 3 | c.2279G>A | TGM1 | Lamellar Ichthyosis | Yes |
|  | Sample 4 | c.2279G>A | TGM1 | Lamellar Ichthyosis | Yes |
|  | Sample 5 | c.2279G>A | TGM1 | Lamellar Ichthyosis | Yes |
|  | Sample 6 | c.2279G>A | TGM1 | Lamellar Ichthyosis | Yes |
|  | Sample 7 | Negative |  |  | No |
|  | Sample 8 | Negative |  |  | No |
|  | Sample 9 | Negative |  |  | No |
| Family 2 | Sample 10 | c.901-1G>A | ABCD1 | X-linked Adrenoleukodystrophy | Yes |
|  | Sample 11 | c.901-1G>A | ABCD1 | X-linked Adrenoleukodystrophy | Yes |
|  | Sample 12 | c.901-1G>A | ABCD1 | X-linked Adrenoleukodystrophy | Yes |
|  | Sample 13 | c.901-1G>A | ABCD1 | X-linked Adrenoleukodystrophy | Yes |
|  | Sample 14 | c.901-1G>A | ABCD1 | X-linked Adrenoleukodystrophy | Yes |
|  | Sample 15 | Negative |  |  | No |
|  | Sample 16 | Negative |  |  | No |
|  | Sample 17 | c.901-1G>A | ABCD1 | X-linked Adrenoleukodystrophy | Yes |
|  | Sample 18 | c.901-1G>A | ABCD1 | X-linked Adrenoleukodystrophy | Yes |
|  | Sample 19 | c.901-1G>A | ABCD1 | X-linked Adrenoleukodystrophy | Yes |
|  | Sample 20 | c.901-1G>A | ABCD1 | X-linked Adrenoleukodystrophy | Yes |
|  | Sample 21 | c.901-1G>A | ABCD1 | X-linked Adrenoleukodystrophy | Yes |

**Table 3.** Variants in Coriell cell lines that possess known variants. PGT-WGS captured all reported variants in positive samples.

| Sample | Variants Reported in Coriell | Gene | Disease | Detected in Orchid WGS? |
| --- | --- | --- | --- | --- |
| NA09301 | NM_003640.5:c.2204+6T>C | ELP1 | Dysautonomia | Yes |
| NA19977 | NM_022455.5:c.6450dup(p.Lys2151GlnfsTer15) | NSD1 | Sotos Syndrome | Yes |
| NA07500 | NM_000038.6:c.4612_4613del(p.Glu1538IlefsTer5) | APC | Familial Adenomatous Polyposis | Yes |
| NA14090 | NM_007300.4:c.68_69del(p.Glu23ValfsTer17) | BRCA1 | Hereditary Breast and Ovarian Cancer | Yes |
| NA14170 | NM_000059.4:c.5946del(p.Ser1982ArgfsTer22) | BRCA2 | Hereditary Breast and Ovarian Cancer | Yes |
| NA13715 | NM_007300.4:c.5329dup(p.Gln1777ProfsTer74) | BRCA1 | Hereditary Breast and Ovarian Cancer | Yes |
| NA00998 | NM_000492.4:c.1521_1523del(p.Phe508del) | CFTR | Cystic Fibrosis | Yes |
| NA13591 | NM_000492.4:c.1521_1523del(p.Phe508del)<br>NM_000492.4:c.350G>A(p.Arg117His)<br>NM_000410.4:c.187C>G(p.His63Asp)<br>NM_005957.5:c.665C>T(p.Ala222Val) | CFTR<br>CFTR<br>HFE<br>MTHFR | Cystic Fibrosis, Hereditary Hemochromatosis, Homocystinuria (MTHFR related) | Yes |
| NA10080 | NM_000314.8:c.781C>T(p.Gln261Ter) | PTEN | PTEN hamartoma tumor syndrome | Yes |
| NA13326 | NM_000051.4:c.1179_1180del(p.Trp393Ter)<br>NM_000051.4:c.4396C>T(p.Arg1466Ter)<br>NM_024675.4:c.3508C>T(p.His1170Tyr)<br>NM_000535.7:c.88C>A(p.Gln30Lys) | ATM<br>ATM<br>PALB2<br>PMS2 | Ataxia Telangiectasia, Lynch Syndrome | Yes |
| NA12932 | NM_001079804.3:c.1441T>C(p.Trp481Arg)<br>NM_001079804.3:c.1326+1G>A | GAA<br>GAA | Glycogen storage disease Type 2 | Yes |
| NA14646 | NM_000410.4:c.845G>A(p.Cys282Tyr) | HFE | Hereditary Hemochromatosis | Yes |
| NA22752 | NM_000090.4:c.636+5G>A | COL3A1 | Ehlers-Danlos syndrome | Yes |
| NA00325 | NM_000402.4:c.653C>T(p.Ser218Phe) | G6PD | G6PD Deficiency | Yes |
| NA12878 | NM_000769.4:c.681G>A(p.Pro227=) | CYP2C19 | Clopidogrel resistance | Yes |
| NA11319 | NM_001127328.2:c.178G>C(p.Ala60Pro) | ACADM | Medium Chain Acyl-CoA Dehydrogenase Deficiency (MCAD) | Yes |
| NA07859 | NM_000492.4:c.3302T>A(p.Met1101Lys) | CFTR | Cystic Fibrosis | Yes |
| NA07415 | NM_000368.5:c.993_994insA(p.Ser332IlefsTer9) | TSC1 | Tuberous Sclerosis Complex | Yes |
| NA21976 | NM_000138.5:c.3444dup(p.Asn1149GlnfsTer10) | FBN1 | Marfan Syndrome | Yes |
| NA11906 | MT:8344 A⇒G |  | MT-TK (myoclonic epilepsy with ragged-red fibers) | Yes |
| NA04368 | ENST00000361381.2:c.1019G>A(p.Arg340His) | MT-ND4 | Leber Hereditary Optic Neuropathy | Yes |
| NA13741 | ENST00000361899.2:c.467T>G(p.Leu156Arg) | MT-ATP6 | Mitochondrial Complex V Deficiency | Yes |
| NA10742 | ENST00000361381.2:c.1019G>A(p.Arg340His) | MT-ND4 | Leber Hereditary Optic Neuropathy | Yes |

We demonstrate the value of this screening via a clinical research case (Table S6). Here, both parents carry different HOGA1 variants as determined by carrier screening. The carrier screening included analysis for pathogenic variants on ∼1300 genes related to neurodevelopmental disorders, birth defects and hereditary cancers based on ACMG panels, as well as the HOGA1 variants identified on previous carrier analysis. Embryo HOGA1 variants were screened from WGS data directly without use of parental genomes. From this batch of embryos we were able to identify a euploid non-carrier embryo available for implantation without parental genotypes or custom probes.

### Common microduplications/deletions, triploid and uniparental disomy screening

Besides variants, we also validated the screening for microduplications, microdeletions, and uniparental disomy etc that conventional PGT-A is unable to detect. Amplified DNA from picogram-level of Coriell cell lines were used as a substitute since availability of research embryos with the above genotypes are extremely rare. We covered a variety of copy number variation related diseases including but not limited to Cri-du-chat, Williams-Beuren, DiGeorge and Prader-Willi syndrome (Table S7). Based on our limit of detection study, we were able to reproducibly detect copy number variation greater than 400 Kb and UPD greater than 20 Mb at a sequence depth of 30X WGS. Examples are shown in Figure S3 and Table S7.

## Discussion

We investigated the performance of a new protocol for amplifying extremely limited amounts of DNA obtained from embryo biopsies. We considered PGT-A, PGT-whole genome sequencing, mitochondrial DNA, microduplication, microdeletions and uniparental disomy using our latest lab developed assay. Remarkably, the DNA genomic coverage, sensitivity, and specificity from amplified material was broadly equivalent to assays performed on the current clinical gold standard, unamplified genomic DNA. Based on equivalent coverage, sensitivity, and specificity, this technique should make it possible to screen embryos before implantation for chromosomal and genetic disorders. Potential benefits of the new technique may include increasing the likelihood of a successful pregnancy (based on the fraction of mitochondrial reads) and reducing the likelihood of birth defects or pediatric or adult developmental disorders.

To ensure accurate and relevant identification of genetic causes of severe, highly penetrant monogenic diseases, we scored 1,300 genes associated with well-studied conditions(9). These gene selections are based on ACMG panels for applicable embryo screening and are grounded in extensive research spanning decades, including familial analysis and cohort studies(10). For many of these genes, functional investigations have established strong links between them and specific disorders. Considering that humans possess over 20,000 genes, our approach takes a conservative stance by concentrating on the 1,300 genes that have undergone comprehensive review and have been unequivocally linked to severe disease. This targeted gene selection ensures that we prioritize clinically meaningful implications for patients when making decisions about embryo transfer. Traditional carrier screening, for example, requires genes to have a confirmed relationship between the detected mutations and a well-defined disorder to provide valuable information to patients to allow reproductive decision making. It is crucial to strike a balance between the amount of data provided and the complexity it presents, as an excess of information can pose challenges when patients are asked to make reproductive decisions. We similarly prioritize diagnostic precision, giving high accuracy in identifying the genetic causes of severe monogenic diseases.

Polygenic risk scores are also easily computable with our method due to high genomic coverage and accuracy in variants. Some polygenic risk scores utilize 6 - 7 million SNPs in total(11), which is difficult to achieve on SNP array chips without relying heavily on statistical imputation. The theoretical limits of genome imputation as applied to rare variant detection is a topic of open research(12).

## Conclusion

To our knowledge, this is the first clinical validation of whole genome embryo screening. In this study, we demonstrated high accuracy for aneuploidy calls (>99.9%) and genetic variants (99.99%), even in the absence of parental genomes. This assay demonstrates advancements in genomic screening and an extended scope for testing capabilities in the realm of preimplantation genetic testing.

## Author Contributions

YX, JS designed experiments. BB, NRS, NS, XD, JK, RB, EU contributed key materials, methods, and discussion. YX, DC, EB, BP, MH, QZ, JK, EB, JC performed and analyzed experiments. YX, WC, MK wrote the manuscript.

## Acknowledgements

We would like to acknowledge HRC Fertility for their assistance in the study especially Bar Sverdlov, Nadia Deratani and Christopher Valdez. We would like to thank Psomagen (CAP & CLIA) for the sequencing service.

---

**Figure S1.**
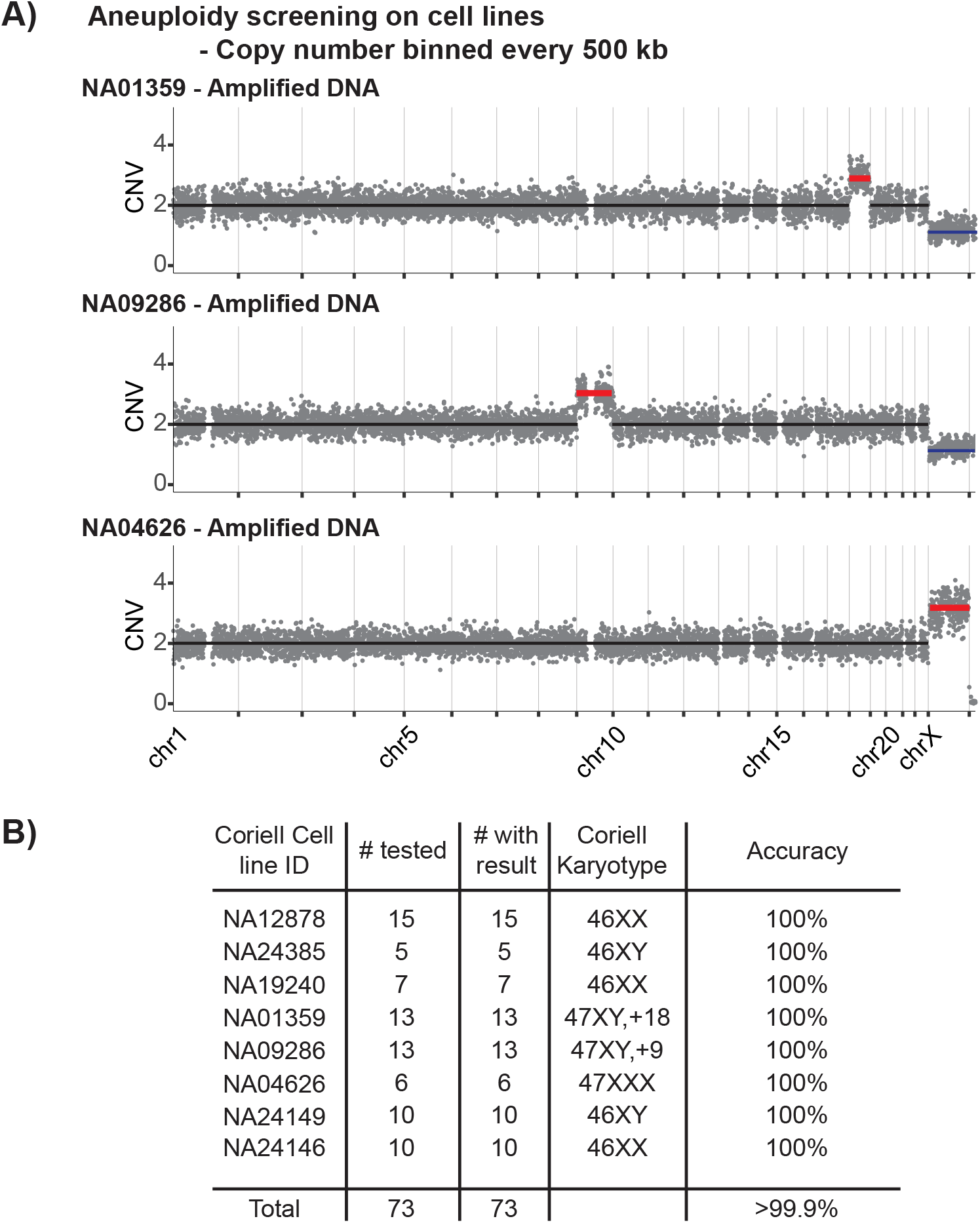
Summary of PGT-A on cell lines. A) Examples of copy number results on cell lines. B) List of all 73 samples with Coriell cell lines.

**Figure S2.**
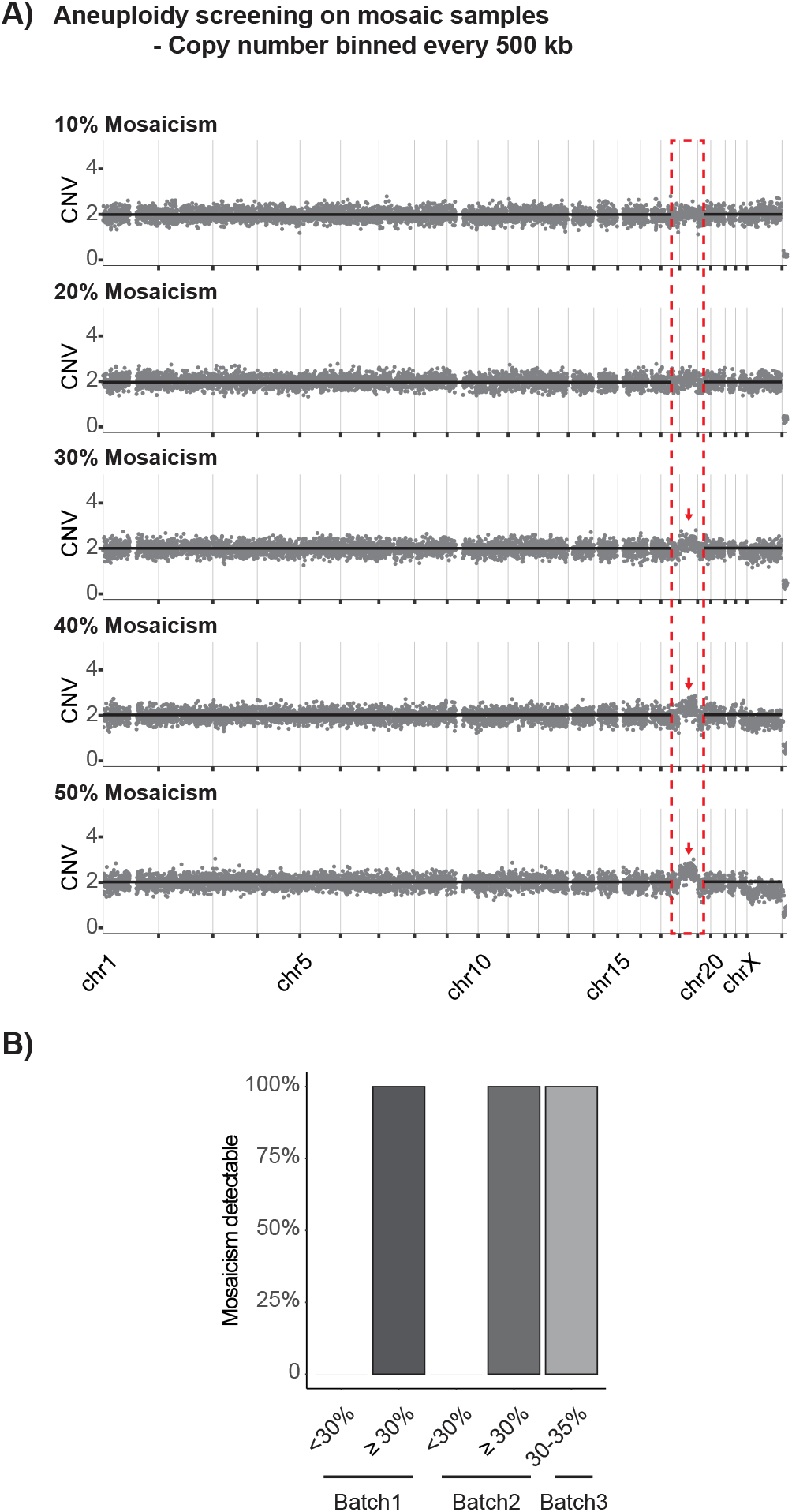
Mosaicism is reported at 30%. A) Mosaicism was reported at 30% level. B) For both batch 1 & 2, no samples were reported at < 30% mosaicism, but >99.9% samples were reported at >=30%. Batch 3 has 24 samples from 30% to 35% mosaicism, and >99.9% were reported.

**Figure S3.**
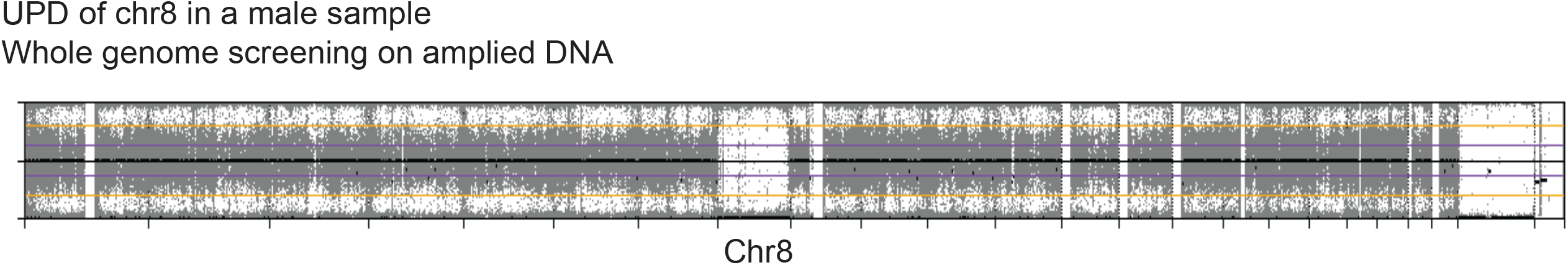
Plot of beta-allele frequency on the whole genome indicates UPD on chromosome 8. Chromosome X and Y had no beta-allele frequency. This is because male has one X and one Y so that beta-allele frequency is zero.

**Table S1.** List of embryo samples in PGT-A validation.

| Sample ID | Original Report Date | Original PGT results | Orchid Collection Date | Orchid Blind Call | Sample ID | Original Report Date | Original PGT results | Orchid Collection Date | Orchid Blind Call |
| --- | --- | --- | --- | --- | --- | --- | --- | --- | --- |
| 1 | 2/20/2016 | XY, +8 | 11/11/2021 | XY, +8 | 56 | 6/22/2017 | XX, -2 | 1/10/2022 | XX, -2p |
| 2 | 2/20/2016 | XY, +8 | 11/11/2021 | XY, +8 | 57 | 6/22/2017 | XX, -2 | 1/10/2022 | XX, -2 |
| 3 | 2/20/2016 | XY, +8 | 11/11/2021 | XY, +8 | 58 | 7/23/2018 | XY, -18, +22 | 2/2/2022 | XY, -18, +22 |
| 4 | 2/20/2016 | XX, +4, -10, -18 | 11/11/2021 | XX, +4, -10, -18 | 59 | 7/23/2018 | XY, -18, +22 | 2/2/2022 | XY, -18, +22 |
| 5 | 2/20/2016 | XX, +4, -10, -18 | 11/11/2021 | XX, +4, -10, -18 | 60 | 12/18/2017 | XX, -2, -14 | 2/2/2022 | XX, -2, -14 |
| 6 | 2/20/2016 | XX, +4, -10, -18 | 11/11/2021 | XX, +4, -10, -18 | 61 | 12/18/2017 | XX, -2, -14 | 2/2/2022 | XX, -2, -14 |
| 7 | 2/20/2016 | XY, +15 | 11/11/2021 | XY, +15 | 62 | 12/18/2017 | XX, -2, -14 | 2/2/2022 | XX, -2, -14 |
| 8 | 2/20/2016 | XY, +15 | 11/11/2021 | XY, +15 | 63 | 12/18/2017 | XY, +5 | 2/2/2022 | XY, +5 |
| 9 | 2/20/2016 | XY, +15 | 11/11/2021 | XY, +15 | 64 | 12/18/2017 | XY, +5 | 2/2/2022 | XY, +5 |
| 10 | 2/20/2016 | XX, -20 | 11/11/2021 | XX, -20 | 65 | 12/18/2017 | XY, +5 | 2/2/2022 | XY, +5 |
| 11 | 2/20/2016 | XX, -20 | 11/11/2021 | Indeterminate | 66 | 12/17/2015 | XX | 2/2/2022 | XX |
| 12 | 2/20/2016 | XX, -20 | 11/11/2021 | XX, -20 | 67 | 12/17/2015 | XX | 2/2/2022 | XX |
| 13 | 2/20/2016 | XY, -13, -16 | 11/11/2021 | XY, -13, -16 | 68 | 12/17/2015 | XX | 2/2/2022 | XX |
| 14 | 2/20/2016 | XY, -13, -16 | 11/11/2021 | XY, -13, -16 | 69 | 12/17/2015 | XX, -16 | 2/2/2022 | XX, -16 |
| 15 | 2/20/2016 | XY, -13, -16 | 11/11/2021 | XY, -13, -16 | 70 | 12/17/2015 | XX, -16 | 2/2/2022 | XX, -16 |
| 16 | 2/20/2016 | XX, -21 | 11/11/2021 | XX, -21 | 71 | 12/17/2015 | XX, -16 | 2/2/2022 | XX, -16 |
| 17 | 2/20/2016 | XX, -21 | 11/11/2021 | XX, -21 | 72 | 6/8/16 | XY, -16 | 2/14/2022 | XY, -16 |
| 18 | 2/20/2016 | XX, -21 | 11/11/2021 | XX, -21 | 73 | 6/8/16 | XY, -16 | 2/14/2022 | XY, -16 |
| 19 | 2/20/2016 | XY, +7, -15 | 11/11/2021 | XY, +7, -15 | 74 | 6/8/16 | XY, -16 | 2/14/2022 | XY, -16 |
| 20 | 2/20/2016 | XY, +7, -15 | 11/11/2021 | XY, +7, -15 | 75 | 10/18/21 | XX | 2/14/2022 | XX |
| 21 | 2/20/2016 | XY, +7, -15 | 11/11/2021 | XY, +7, -15 | 76 | 10/19/21 | XX | 2/14/2022 | XX |
| 22 | 7/4/2013 | XY | 12/6/2021 | XY | 77 | 10/20/21 | XX | 2/14/2022 | XX |
| 23 | 7/4/2013 | XY | 12/6/2021 | XY | 78 | 11/9/2018 | 46, No sex reported | 8/10/2022 | XX |
| 24 | 7/4/2013 | XY | 12/6/2021 | XY | 79 | 11/9/2018 | 46, No sex reported | 8/10/2022 | XX |
| 25 | 7/4/2013 | XX | 12/6/2021 | XX | 80 | 11/9/2018 | 46, No sex reported | 8/10/2022 | XX |
| 26 | 7/4/2013 | XX | 12/6/2021 | Indeterminate | 81 | 11/9/2018 | 46, No sex reported | 8/10/2022 | XY |
| 27 | 7/4/2013 | XX | 12/6/2021 | XX | 82 | 11/9/2018 | 46, No sex reported | 8/10/2022 | XY |
| 28 | 7/4/2013 | XY, -5, +18 | 12/6/2021 | XY, -5 | 83 | 11/9/2018 | 46, No sex reported | 8/10/2022 | XY |
| 29 | 7/4/2013 | XY, -5, +18 | 12/6/2021 | XY, -5 | 84 | 11/9/2018 | 46, No sex reported | 8/10/2022 | XX |
| 30 | 7/4/2013 | XY, -5, +18 | 12/6/2021 | XY, -5 | 85 | 11/9/2018 | 46, No sex reported | 8/10/2022 | XX |
| 31 | 7/4/2013 | XY, +2, +22 | 12/6/2021 | XY, -2, +22 | 86 | 11/9/2018 | 46, No sex reported | 8/10/2022 | XX |
| 32 | 7/4/2013 | XY, +2, +22 | 12/6/2021 | XY, -2, +22 | 87 | 11/9/2018 | 46, No sex reported | 8/10/2022 | XX |
| 33 | 7/4/2013 | XY, +2, +22 | 12/6/2021 | XY, -2, +22 | 88 | 11/9/2018 | 46, No sex reported | 8/10/2022 | XX |
| 34 | 7/4/2013 | XX, +9, +13, +21 | 12/6/2021 | XY, +9, +13, +21 | 89 | 11/9/2018 | 46, No sex reported | 8/10/2022 | XX |
| 35 | 7/4/2013 | XX, +9, +13, +21 | 12/6/2021 | XY, +9, +13, +21 | 90 | 11/9/2018 | 46, No sex reported | 8/10/2022 | XY |
| 36 | 7/4/2013 | XX, +9, +13, +21 | 12/6/2021 | XY, +9, +13, +21 | 91 | 11/9/2018 | 46, No sex reported | 8/10/2022 | XY |
| 37 | 7/4/2013 | XY | 12/6/2021 | XY | 92 | 11/9/2018 | 46, No sex reported | 8/10/2022 | XY |
| 38 | 7/4/2013 | XY | 12/6/2021 | XY | 93 | 11/9/2018 | 46, No sex reported | 8/10/2022 | XX |
| 39 | 7/4/2013 | XY | 12/6/2021 | XY | 94 | 11/9/2018 | 46, No sex reported | 8/10/2022 | XX |
| 40 | 9/26/2016 | XX | 12/6/2021 | XX | 95 | 11/9/2018 | 46, No sex reported | 8/10/2022 | XX |
| 41 | 9/26/2016 | XX | 12/6/2021 | XX | 96 | 11/17/2017 | XY | 8/10/2022 | XY |
| 42 | 9/26/2016 | XX | 12/6/2021 | XX | 97 | 11/17/2017 | XY | 8/10/2022 | XY |
| 43 | 10/5/2020 | XX, -3, +15, +20, +21, +22 | 12/6/2021 | XX, -3, +15, +20, 21, 22 | 98 | 11/17/2017 | XY | 8/10/2022 | XY |
| 44 | 10/5/2020 | XX, -3, +15, +20, +21, +22 | 12/6/2021 | XX, -3, +15, +20, 21, 22 | 99 | 11/17/2017 | XY | 8/10/2022 | XY |
| 45 | 10/5/2020 | XX, -3, +15, +20, +21, +22 | 12/6/2021 | XX, -3, +15, +20, 21, 22 | 100 | 11/17/2017 | XY | 8/10/2022 | XY |
| 46 | 4/25/2017 | XY | 1/10/2022 | XY | 101 | 11/17/2017 | XY | 8/10/2022 | XY |
| 47 | 4/25/2017 | XY | 1/10/2022 | XY | 102 | 11/17/2017 | XX, +22 | 8/10/2022 | XX, +22 |
| 48 | 4/25/2017 | XY | 1/10/2022 | XY | 103 | 11/17/2017 | XX, +22 | 8/10/2022 | XX, +22 |
| 49 | 4/25/2017 | XY | 1/10/2022 | XY | 104 | 9/6/2017 | XX, +19 | 8/10/2022 | XX, +19 mosaic |
| 50 | 4/25/2017 | XY | 1/10/2022 | XY | 105 | 9/6/2017 | XX, +19 | 8/10/2022 | XX, +19 |
| 51 | 4/25/2017 | XY | 1/10/2022 | XY | 106 | 10/12/2022 | XX, -18 | 12/8/2022 | XX, -18 |
| 52 | 6/22/2017 | XX, -16 | 1/10/2022 | XX, -16 | 107 | 10/12/2022 | XX, -18 | 12/8/2022 | XX, -18 |
| 53 | 6/22/2017 | XX, -16 | 1/10/2022 | XX, -16 | 108 | 10/12/2022 | XX, -18 | 12/8/2022 | XX, -18 |
| 54 | 6/22/2017 | XX, -16 | 1/10/2022 | XX, -16 | 109 | 12/8/2020 | XY | 12/8/2022 | XY |
| 55 | 6/22/2017 | XX, -2 | 1/10/2022 | XX, -2p | 110 | 12/8/2020 | XY | 12/8/2022 | XY |

**Table S2.** Comparison of gDNA and amplified DNA from NA12878 in terms of genomic coverage, total SNV called, accuracy, specificity, precision, and sensitivity. Variants list of NA12878 from NIST was used as the reference.

| Sample | Genomic Coverage | Total SNV | Accuracy | Specificity | Precision | Sensitivity |
| --- | --- | --- | --- | --- | --- | --- |
| NA12878 gDNA Sample 1 | 99.8% | 3286979 | 99.996% | 99.997% | 97.8% | 98.8% |
| NA12878 gDNA Sample 2 | 99.9% | 3290927 | 99.996% | 99.997% | 97.8% | 99.0% |
| NA12878 gDNA Sample 3 | 99.8% | 3286497 | 99.996% | 99.997% | 97.8% | 98.8% |
| NA12878 gDNA Sample 4 | 99.9% | 3287953 | 99.996% | 99.997% | 97.9% | 99.0% |
| NA12878 gDNA Sample 5 | 99.8% | 3277286 | 99.996% | 99.997% | 98.0% | 98.8% |
| **Average** | **99.8%** | **3285928** | **99.996%** | **99.997%** | **97.8%** | **98.9%** |
| NA12878 Amplified DNA Sample 1 | 99.7% | 3238978 | 99.994% | 99.997% | 97.9% | 97.6% |
| NA12878 Amplified DNA Sample 2 | 99.8% | 3263126 | 99.995% | 99.997% | 98.0% | 98.4% |
| NA12878 Amplified DNA Sample 3 | 99.8% | 3252094 | 99.995% | 99.998% | 98.2% | 98.3% |
| NA12878 Amplified DNA Sample 4 | 99.8% | 3258943 | 99.996% | 99.998% | 98.2% | 98.4% |
| NA12878 Amplified DNA Sample 5 | 99.7% | 3229761 | 99.994% | 99.997% | 97.9% | 97.3% |
| **Average** | **99.8%** | **3248580** | **99.995%** | **99.997%** | **98.1%** | **98.0%** |

**Table S3.** Tabulated numbers for true positive, false positive, true negative and false negative counts from NA12878 samples. These numbers were used to compute Table S2.

| Sample | True Positives | False Positives | True Negatives | False Negatives |
| --- | --- | --- | --- | --- |
| NA12878 gDNA Sample 1 | 3214156 | 73257 | 2498498635 | 37667 |
| NA12878 gDNA Sample 2 | 3218277 | 73072 | 2498498820 | 33546 |
| NA12878 gDNA Sample 3 | 3214192 | 72753 | 2498499139 | 37631 |
| NA12878 gDNA Sample 4 | 3218242 | 70121 | 2498501771 | 33581 |
| NA12878 gDNA Sample 5 | 3212093 | 65599 | 2498506293 | 39730 |
| **Average** | **3215392** | **70960** | **2498500932** | **36431** |
| NA12878 Amplified DNA Sample 1 | 3172496 | 67132 | 2498504760 | 79327 |
| NA12878 Amplified DNA Sample 2 | 3199885 | 63958 | 2498507934 | 51938 |
| NA12878 Amplified DNA Sample 3 | 3194879 | 57880 | 2498514012 | 56944 |
| NA12878 Amplified DNA Sample 4 | 3200575 | 59098 | 2498512794 | 51248 |
| NA12878 Amplified DNA Sample 5 | 3162468 | 67937 | 2498503955 | 89355 |
| **Average** | **3186060** | **63201** | **2498508691** | **65763** |

**Table S4.** Tabulated numbers for true positive, false positive, true negative and false negative counts from embryo samples. These numbers were used to compute Table 1.

| Sample | True Positives | False Positives | True Negatives | False Negatives |
| --- | --- | --- | --- | --- |
| 1 | 3270826 | 72978 | 2498427158 | 52753 |
| 2 | 3260902 | 66729 | 2498433445 | 62639 |
| 3 | 3241275 | 70864 | 2498447239 | 64337 |
| 4 | 3208229 | 71239 | 2498446853 | 97394 |
| 5 | 3348398 | 55757 | 2498354672 | 64888 |
| 6 | 3358076 | 56201 | 2498354258 | 55180 |
| 7 | 3307540 | 53142 | 2498399224 | 63809 |
| 8 | 3106492 | 57099 | 2498602671 | 57453 |
| 9 | 3105403 | 54710 | 2498605123 | 58479 |
| 10 | 3213236 | 91074 | 2498453268 | 66137 |
| 11 | 3218889 | 82827 | 2498461470 | 60529 |
| 12 | 3322745 | 46089 | 2498393320 | 61561 |
| 13 | 3299943 | 40540 | 2498398825 | 84407 |
| 14 | 3187551 | 83342 | 2498499668 | 53154 |
| 15 | 3184474 | 78092 | 2498504866 | 56283 |
| 16 | 3201676 | 72658 | 2498492003 | 57378 |
| 17 | 3201352 | 73875 | 2498490706 | 57782 |
| 18 | 3192477 | 61970 | 2498516295 | 52973 |
| 19 | 3175772 | 58945 | 2498519275 | 69723 |
| 20 | 3205470 | 68532 | 2498499144 | 50569 |
| 21 | 3196791 | 60570 | 2498507126 | 59228 |
| 22 | 3212393 | 58179 | 2498482967 | 70176 |
| 23 | 3225625 | 59208 | 2498481928 | 56954 |
| 24 | 3169496 | 68217 | 2498487962 | 98040 |
| 25 | 3324367 | 58178 | 2498394241 | 46929 |
| **Average** | **3229576** | **64841** | **2498466148** | **63150** |

**Table S5.** Analysis of mitochondrial variants from PGT-WGS data. Mitochondrial variants from the whole embryos were used as references.

| Sample | Total Bases | True Positive | True Negative | False Positive | False Negative | Accuracy | Specificity | Sensitivity | Precision |
| --- | --- | --- | --- | --- | --- | --- | --- | --- | --- |
| 1 | 16569 | 19 | 16550 | 0 | 0 | 100.000% | 100.0% | 100.0% | 100.0% |
| 2 | 16569 | 19 | 16550 | 0 | 0 | 100.000% | 100.0% | 100.0% | 100.0% |
| 3 | 16569 | 21 | 16548 | 0 | 0 | 100.000% | 100.0% | 100.0% | 100.0% |
| 4 | 16569 | 21 | 16548 | 0 | 0 | 100.000% | 100.0% | 100.0% | 100.0% |
| 5 | 16569 | 32 | 16537 | 0 | 0 | 100.000% | 100.0% | 100.0% | 100.0% |
| 6 | 16569 | 32 | 16537 | 0 | 0 | 100.000% | 100.0% | 100.0% | 100.0% |
| 7 | 16569 | 32 | 16537 | 0 | 0 | 100.000% | 100.0% | 100.0% | 100.0% |
| 8 | 16569 | 25 | 16544 | 0 | 0 | 100.000% | 100.0% | 100.0% | 100.0% |
| 9 | 16569 | 21 | 16548 | 0 | 4 | 99.976% | 100.0% | 84.0% | 100.0% |
| 10 | 16569 | 25 | 16544 | 0 | 0 | 100.000% | 100.0% | 100.0% | 100.0% |
| 11 | 16569 | 25 | 16544 | 0 | 0 | 100.000% | 100.0% | 100.0% | 100.0% |
| 12 | 16569 | 51 | 16518 | 0 | 0 | 100.000% | 100.0% | 100.0% | 100.0% |
| 13 | 16569 | 50 | 16519 | 0 | 1 | 99.994% | 100.0% | 98.0% | 100.0% |
| 14 | 16569 | 6 | 16563 | 0 | 0 | 100.000% | 100.0% | 100.0% | 100.0% |
| 15 | 16569 | 6 | 16563 | 0 | 0 | 100.000% | 100.0% | 100.0% | 100.0% |
| 16 | 16569 | 6 | 16563 | 0 | 0 | 100.000% | 100.0% | 100.0% | 100.0% |
| 17 | 16569 | 6 | 16563 | 0 | 0 | 100.000% | 100.0% | 100.0% | 100.0% |
| 18 | 16569 | 6 | 16563 | 0 | 0 | 100.000% | 100.0% | 100.0% | 100.0% |
| 19 | 16569 | 6 | 16563 | 0 | 0 | 100.000% | 100.0% | 100.0% | 100.0% |
| 20 | 16569 | 25 | 16544 | 0 | 0 | 100.000% | 100.0% | 100.0% | 100.0% |
| 21 | 16569 | 25 | 16544 | 0 | 0 | 100.000% | 100.0% | 100.0% | 100.0% |
| 22 | 16569 | 24 | 16545 | 0 | 1 | 99.994% | 100.0% | 96.0% | 100.0% |
| 23 | 16569 | 25 | 16544 | 0 | 0 | 100.000% | 100.0% | 100.0% | 100.0% |
| 24 | 16569 | 29 | 16540 | 0 | 1 | 99.994% | 100.0% | 96.7% | 100.0% |
| 25 | 16569 | 32 | 16537 | 0 | 0 | 100.000% | 100.0% | 100.0% | 100.0% |
| Average | 16569 | 22.09 | 16,546.91 | 0.00 | 0.26 | 99.998% | >99.99% | 99.0% | >99.99% |

**Table S6.** Biopsy screening results on HOGA1 variants.

| Sample | Chromosome Abornalities | Father Carrier:<br>HOGA1 c.700+5G>T | Mother Carrier:<br>HOGA1 c.110G>T | Genetic disease status | Available for Transfer |
| --- | --- | --- | --- | --- | --- |
| Sample 1 | Not detected | Positive | Positive | Compound Het. | No |
| Sample 2 | Not detected | Positive | Positive | Compound Het. | No |
| Sample 3 | 6p25.3p22.3(1-20560000)X3 | Positive | Negative | Carrier | No |
| Sample 4 | Not detected | Negative | Negative | Non-carrier | Yes |

**Table S7.** Validation on additional chromosomal abnormalities including microduplication, microdeletion, triploid and uniparental disomy.

| Sample | Coriell Reports | Category | Size | Disease | Detectable in Orchid WGS? |
| --- | --- | --- | --- | --- | --- |
| NA05875 | 16p12.1p11.2(27,203,030-32,180,964)x1 | Segmental | 5Mb |  | Yes |
| NA06228 | 11q23.3q25(116188703-134449982)x3<br>22q11.1q11.21(14435170-18676540)x3 | Segmental | 19Mb<br>4Mb |  | Yes<br>Yes |
| NA07736 | 11p14.1p13(28229986-34925688)x1 | Segmental | 7Mb |  | Yes |
| NA10013 | 69,XXX | Triploid | >20Mb |  | Yes |
| NA10995 | 1q31.3q44(192686117-247190999)x3 | Segmental | >20Mb |  | Yes |
| NA14043 | 2p25.3p25.1(5702-9484791)x2 hmz<br>2q22.3q32.3(144864227-195289707)x2 hmz<br>12q14.1q14.3(56656199-63469126)x2 hmz | UPD | >20Mb |  | Yes<br>Yes<br>Yes |
| NA14117 | 5p15.33p14.3(68519-22367289)x1 | Segmental | 20Mb | Cri du chat syndrome | Yes |
| NA14184 | 7q11.23(72297530-73780028)x1 | Segmental | 1.5Mb | Williams-Beuren syndrome | Yes |
| NA15603 | 8p23.3p23.1(161,221-6,815,328)hmz<br>8p23.1p11.21(8,087,143-42,858,299)hmz | UPD | >20Mb |  | Yes |
| NA17942 | 22q11.21(18748427-21611337)x1 | Segmental | 2Mb | DiGeorge syndrome | Yes |
| NA20201 | 15q11.1q13.3(18276328-30630402)x4 | Segmental | 10Mb |  | Yes |
| NA20408 | Prader-Willi syndrome (Region not specified) | UPD | >20Mb | Prader-Willi syndrome | Yes |
| NA22397 | 15q11.2q13.1(21245026-26500067)x3 | Segmental | 6Mb |  | Yes |
| NA22569 | 1p36.33p36.31(615320-6255256)x1 | Segmental | 6Mb | 1p36 deletion syndrome | Yes |

## Reference

1. Vermeesch JR, Voet T, Devriendt K. Prenatal and pre-implantation genetic diagnosis. Nat Rev Genet 2016;17(10):643–56.

2. Sacchi L, Albani E, Cesana A, Smeraldi A, Parini V, Fabiani M, et al. Preimplantation Genetic Testing for Aneuploidy Improves Clinical, Gestational, and Neonatal Outcomes in Advanced Maternal Age Patients Without Compromising Cumulative Live-Birth Rate. J Assist Reprod Genet 2019;36(12):2493–504.

3. Zheng Y, Wang N, Li L, Jin F. Whole genome amplification in preimplantation genetic diagnosis. J Zhejiang Univ Sci B 2011;12(1):1–11.

4. Van der Auwera GA, Carneiro MO, Hartl C, Poplin R, del Angel G, Levy-Moonshine A, et al. From FastQ Data to High-Confidence Variant Calls: The Genome Analysis Toolkit Best Practices Pipeline. Curr Protoc Bioinformatics 2013;43(1):11.10.1–11.10.33.

5. Dickson D, Tao T, Qin W. Exploring mitochondrial DNA content as a novel biomarker to improve embryo implantation potential: A review. Biomedical Technology 2024;5:82–6.

6. Abhari S, Kawwass JF. Pregnancy and Neonatal Outcomes after Transfer of Mosaic Embryos: A Review. J Clin Med 2021;10(7).

7. Treff NR, Zimmerman R, Bechor E, Hsu J, Rana B, Jensen J, et al. Validation of concurrent preimplantation genetic testing for polygenic and monogenic disorders, structural rearrangements, and whole and segmental chromosome aneuploidy with a single universal platform. Eur J Med Genet 2019;62(8):103647.

8. Huang L, Ma F, Chapman A, Lu S, Xie XS. Single-Cell Whole-Genome Amplification and Sequencing: Methodology and Applications. Annu Rev Genomics Hum Genet 2015;16(1):79–102.

9. Richards S, Aziz N, Bale S, Bick D, Das S, Gastier-Foster J, et al. Standards and guidelines for the interpretation of sequence variants: a joint consensus recommendation of the American College of Medical Genetics and Genomics and the Association for Molecular Pathology. Genetics in Medicine 2015;17(5):405–23.

10. Miller DT, Lee K, Chung WK, Gordon AS, Herman GE, Klein TE, et al. ACMG SF v3.0 list for reporting of secondary findings in clinical exome and genome sequencing: a policy statement of the American College of Medical Genetics and Genomics (ACMG). Genetics in Medicine 2021;23(8):1381–90.

11. Khera A V, Chaffin M, Aragam KG, Haas ME, Roselli C, Choi SH, et al. Genome-wide polygenic scores for common diseases identify individuals with risk equivalent to monogenic mutations. Nat Genet 2018;50(9):1219–24.

12. Si Y, Vanderwerff B, Zöllner S. Why are rare variants hard to impute? Coalescent models reveal theoretical limits in existing algorithms. Genetics 2021;217(4):iyab011.

